## Supplemental Figures for "GlyContact reveals structure-function relationships in glycans"

[*]Corresponding author

### Supplementary Figures

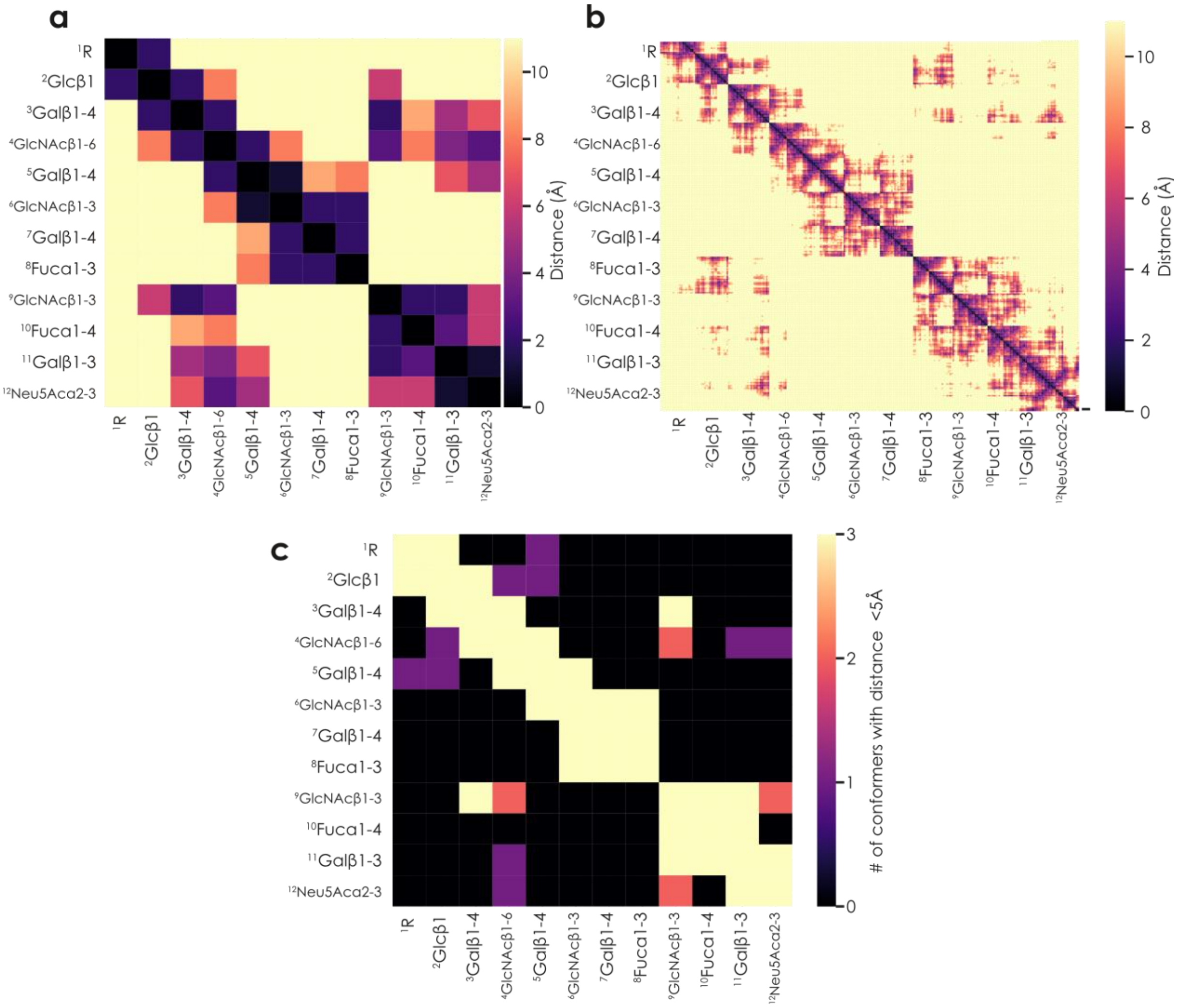


**Supplementary Figure 1. Contact maps for the glycan Fucα1-3(Galβ1-4)GlcNAcβ1-3Galβ1-4GlcNAcβ1-6(Neu5Acα2-3Galβ1-3(Fucα1-4)GlcNAcβ1-3)Galβ1-4Glc. a-c)** For the conformer beta_2 (a-b) or all conformers (c), we calculated their monosaccharide- (a, c) or atom-level (b) contact maps, using the *glycontact.process.make_monosaccharide_contact_table* and *glycontact.process.make_atom_contact_table* functions from GlyContact, respectively. For (c), we then used the *glycontact.process.inter_structure_frequency_table* function to analyze in how many conformers two monosaccharides were spatially adjacent.

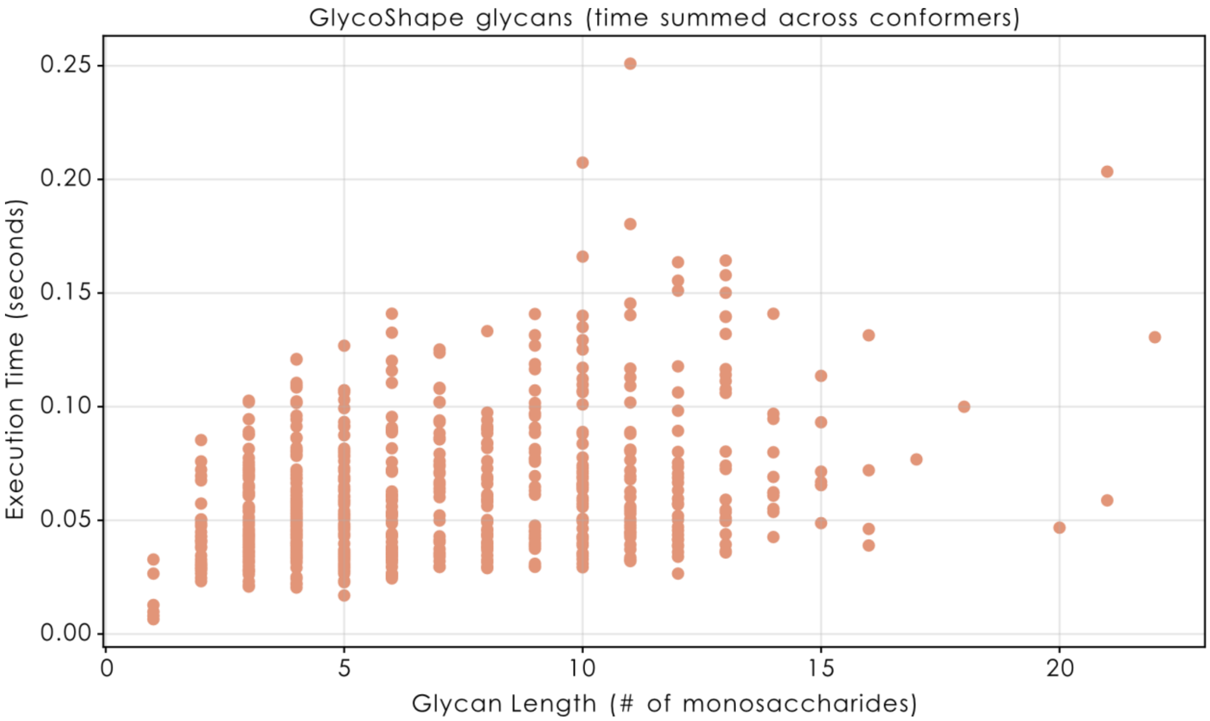


**Supplementary Figure 2. Efficient processing of glycan PDBs with GlyContact.** For all 717 glycans with deposited PDBs on GlycoShape, we processed all their conformers via the *glycontact.process.annotation_pipeline* function, timed the execution, and show the execution time (summed across conformers) for all glycans. Glycan length is calculated by number of monosaccharides.

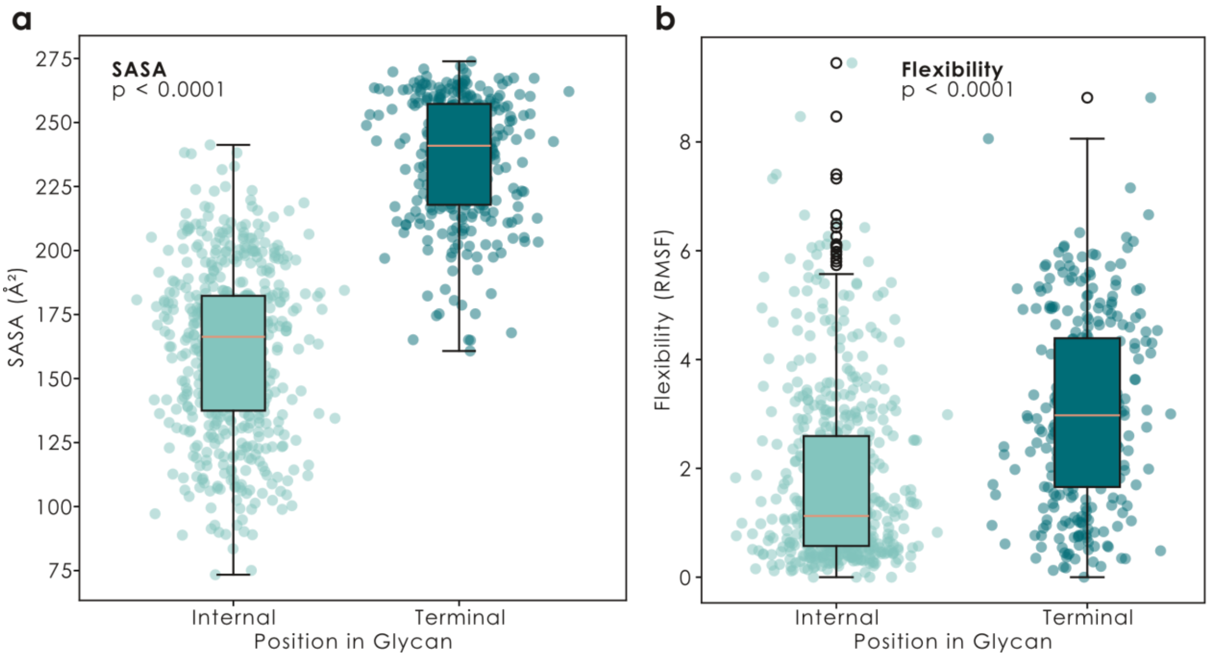


**Supplementary Figure 3. Terminal monosaccharides are more accessible and more flexible. a-b)** For the monosaccharide galactose, we compared the SASA (a) or flexibility (b) values of all occurrences of galactose, across all GlycoShape structures (averaged across conformers, weighted by their proportions), and compare galactose in terminal positions (i.e., non-reducing ends) with internal positions. The data are shown as box plots (line indicating the median, box edges indicating the $25^{th}$ and $75^{th}$ percentile, whiskers indicating the 95% confidence interval, and black circles indicating outliers), as well as an overlaid scatter plot of the actual values, with added horizontal jitter for visibility. Statistical significance was established with a Mann-Whitney U test.

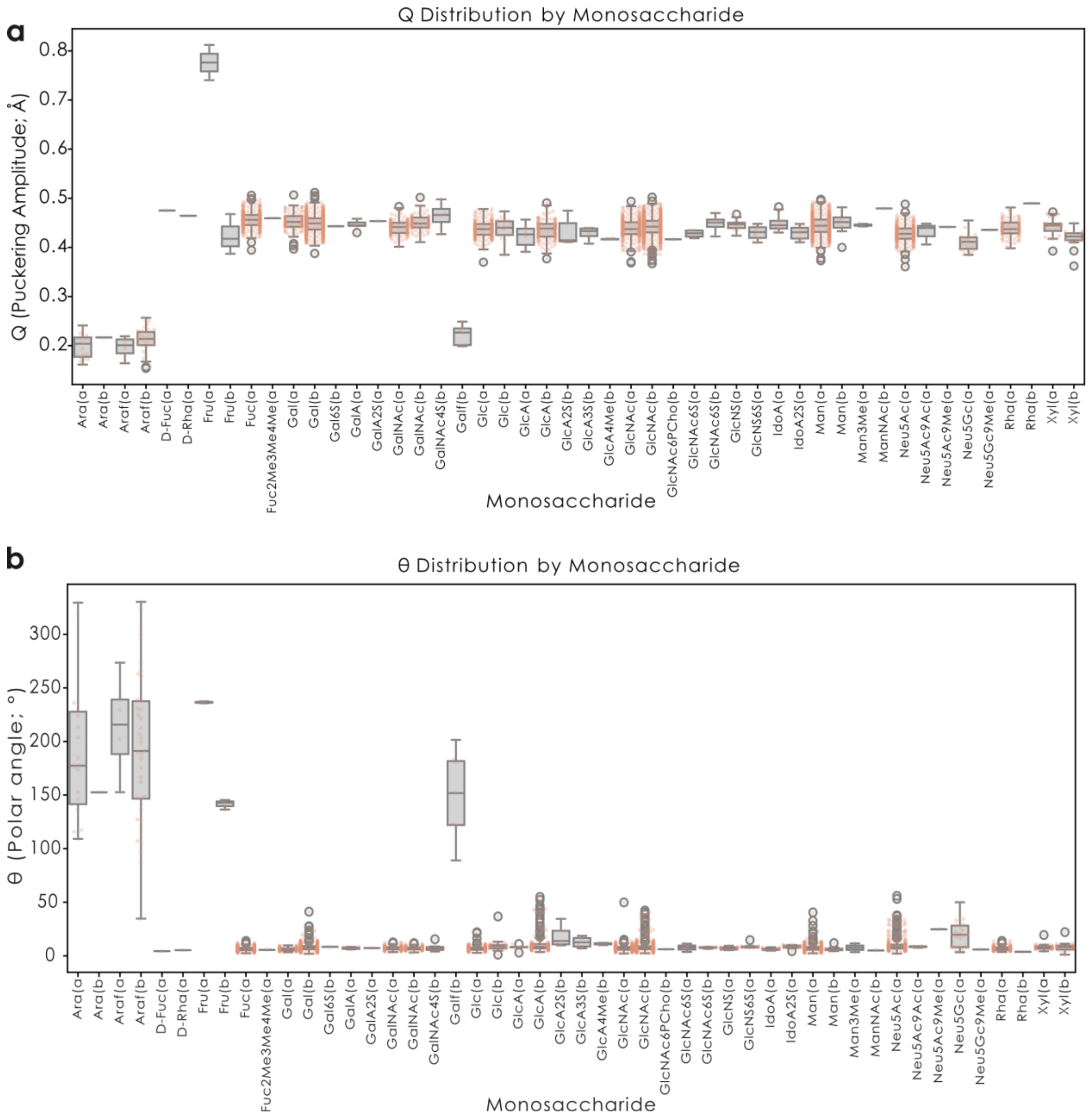


**Supplementary Figure 4. Ring puckering shows similar ranges across monosaccharides. a-b)** For all glycan structures on GlycoShape, we calculated all puckering amplitudes (Q, a) and polar angles (θ, b) and plotted them here per monosaccharide. The data are shown as box plots (line indicating the median, box edges indicating the 25th and 75th percentile, whiskers indicating the 95% confidence interval, and gray circles indicating outliers), as well as an overlaid scatter plot of the actual values, with added horizontal jitter for visibility.

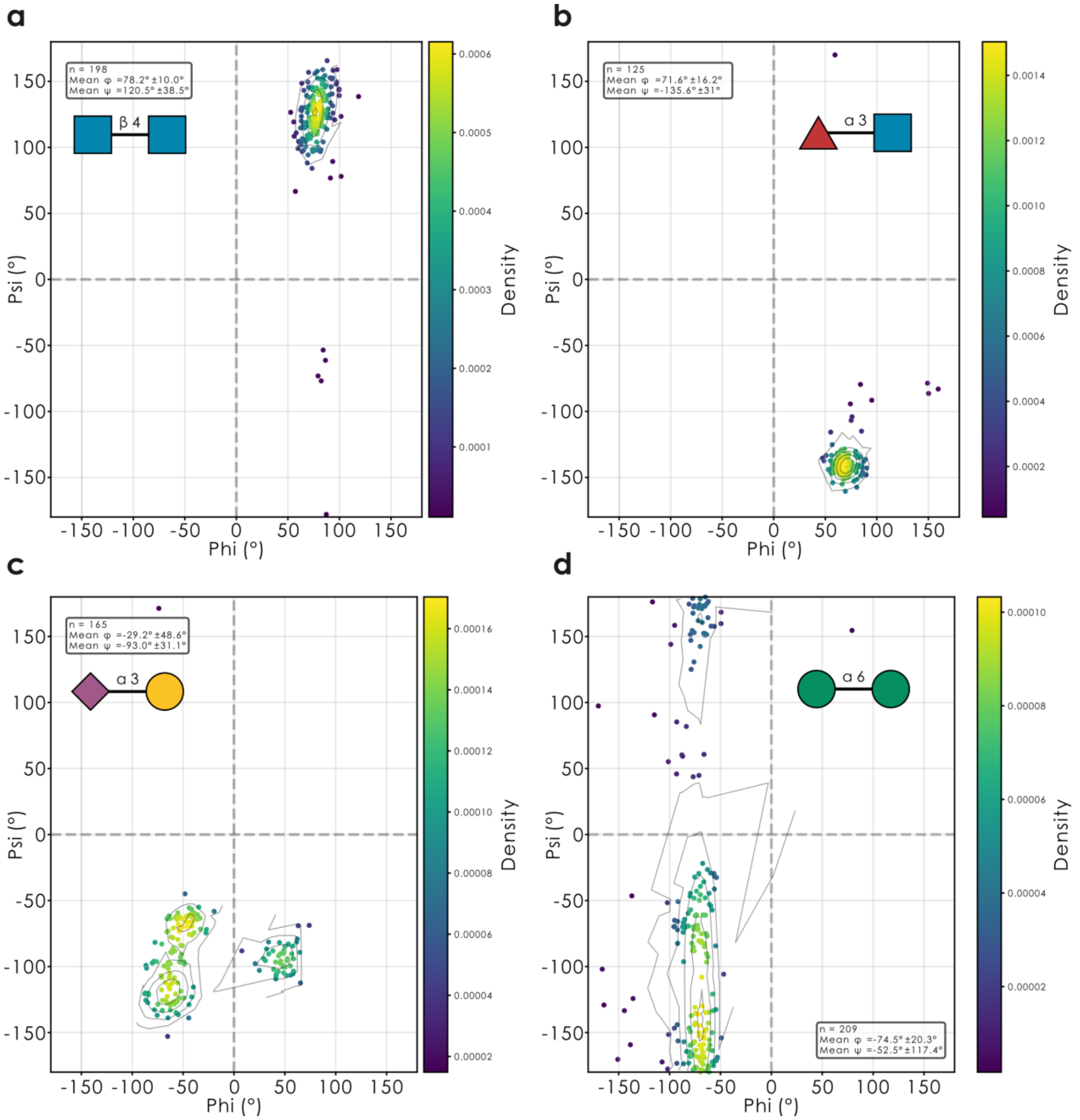


**Supplementary Figure 5. GlyContact-calculated torsion angles conform to common Ramachandran distributions. a-d)** For the example disaccharides GlcNAcβ1-4GlcNAc (a), Fucα1-3GlcNAc (b), Neu5Acα2-3Gal (c), and Manα1-6Man (d), we extracted all their occurrences from all glycan structures available on GlycoShape and calculated their torsion angles (φ, ψ) via the *glycontact.process.get_glycosidic_torsions* function within GlyContact, which is shown here via Ramachandran plots with the contours derived from a Gaussian kernel density estimate. Both number of occurrences and angle mean values are depicted in the respective panel. The entire workflow is available via the *glycontact.visualize.ramachandran_plot* function.

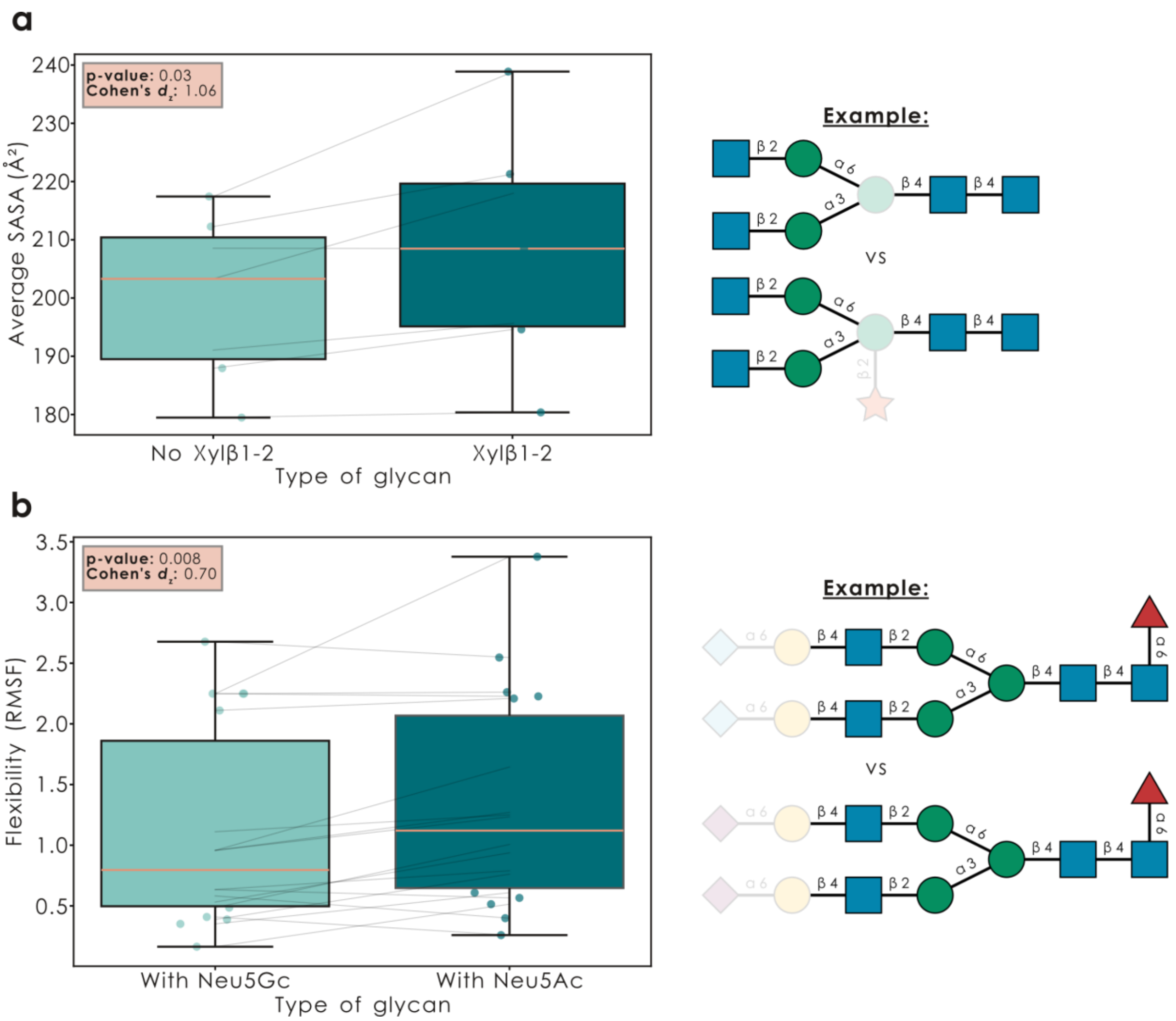


**Supplementary Figure 6. Xylose addition opens up plant *N*-glycans and Neu5Ac increases glycan flexibility. a-b)** For the example of Xylβ1-2 occurrence in *N*-glycans, typical in plants, (a) and Neu5Ac vs Neu5Gc in all GlycoShape glycans (b), we gathered all 'twins' (pairs of sequences that only differed in the presence/absence of Xylβ1-2 or in Neu5Ac vs Neu5Gc, respectively) for which we had structural data from GlycoShape (a, n = 7; b, n = 19) and compared their average SASA (a) or flexibility (b) values (excluding the considered motif and its attachment site). Results are shown as box plots (line indicating the median, box edges indicating the 25$^{th}$ and 75$^{th}$ percentile, and whiskers indicating the 95% confidence interval), as well as an overlaid scatter plot of the actual values, with added horizontal jitter for visibility. Statistical testing involved a paired t-test and Cohen's $d_z$ as an effect size for paired samples. The entire workflow is available via the *glycontact.visualize.find_difference* function.

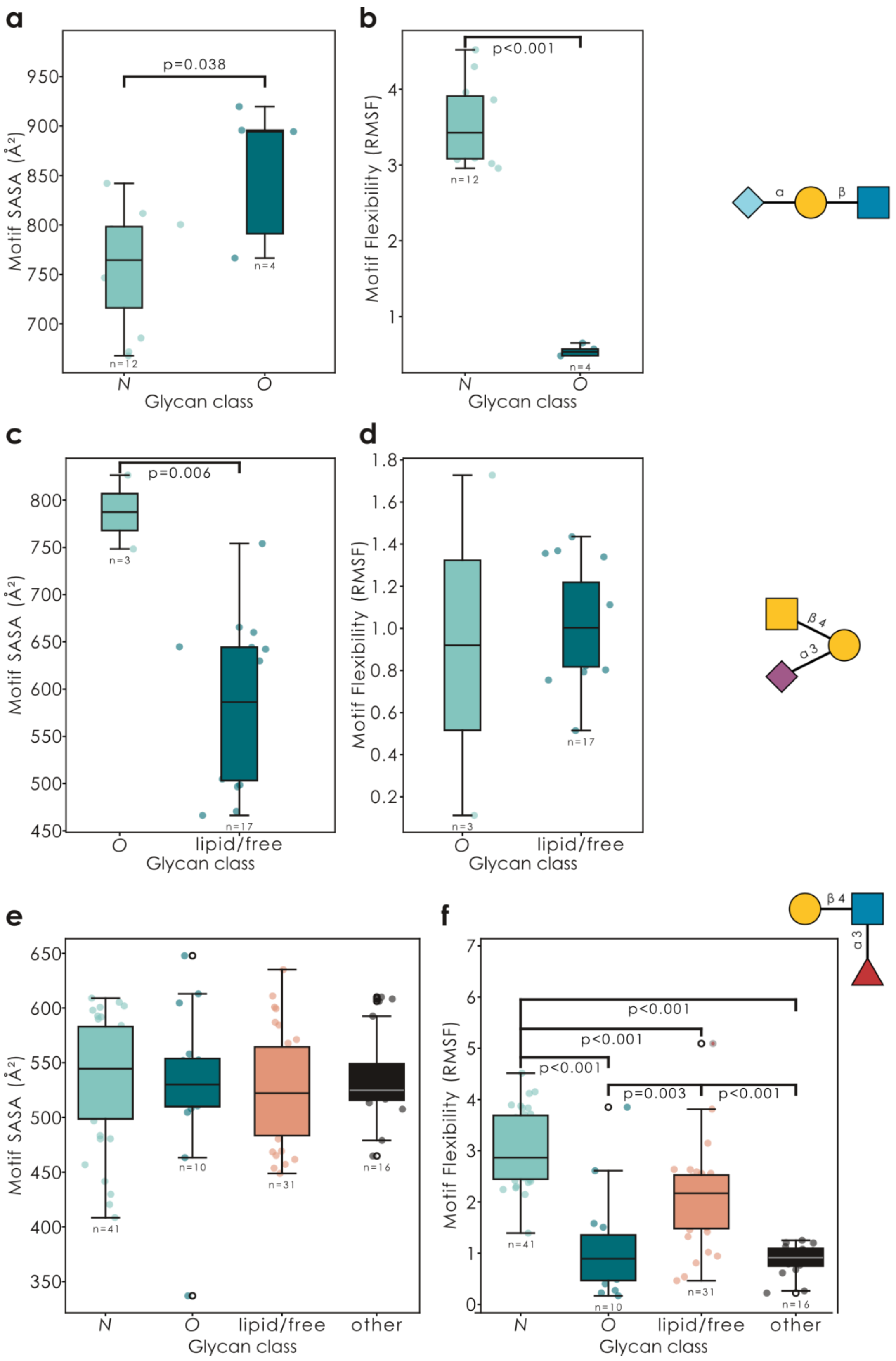


**Supplementary Figure 7. Glycan motifs exposed by different glycan classes exhibit different structural characteristics. a-f)** For Neu5Gc-LacNAc (Neu5Gcα2-?Galβ1-?GlcNAc; a-b), Sd[a] (c-d), and Lewis X (e-f) in GlycoShape glycans, we summed monosaccharide-level SASA values for each motif (a, c, e) and averaged their flexibility (b, d, f). Then, we grouped glycans by glycan class and analyzed differences across classes by an ANOVA, followed by Tukey's HSD post-hoc test. The number of analyzed motif instances is provided under each box plot.

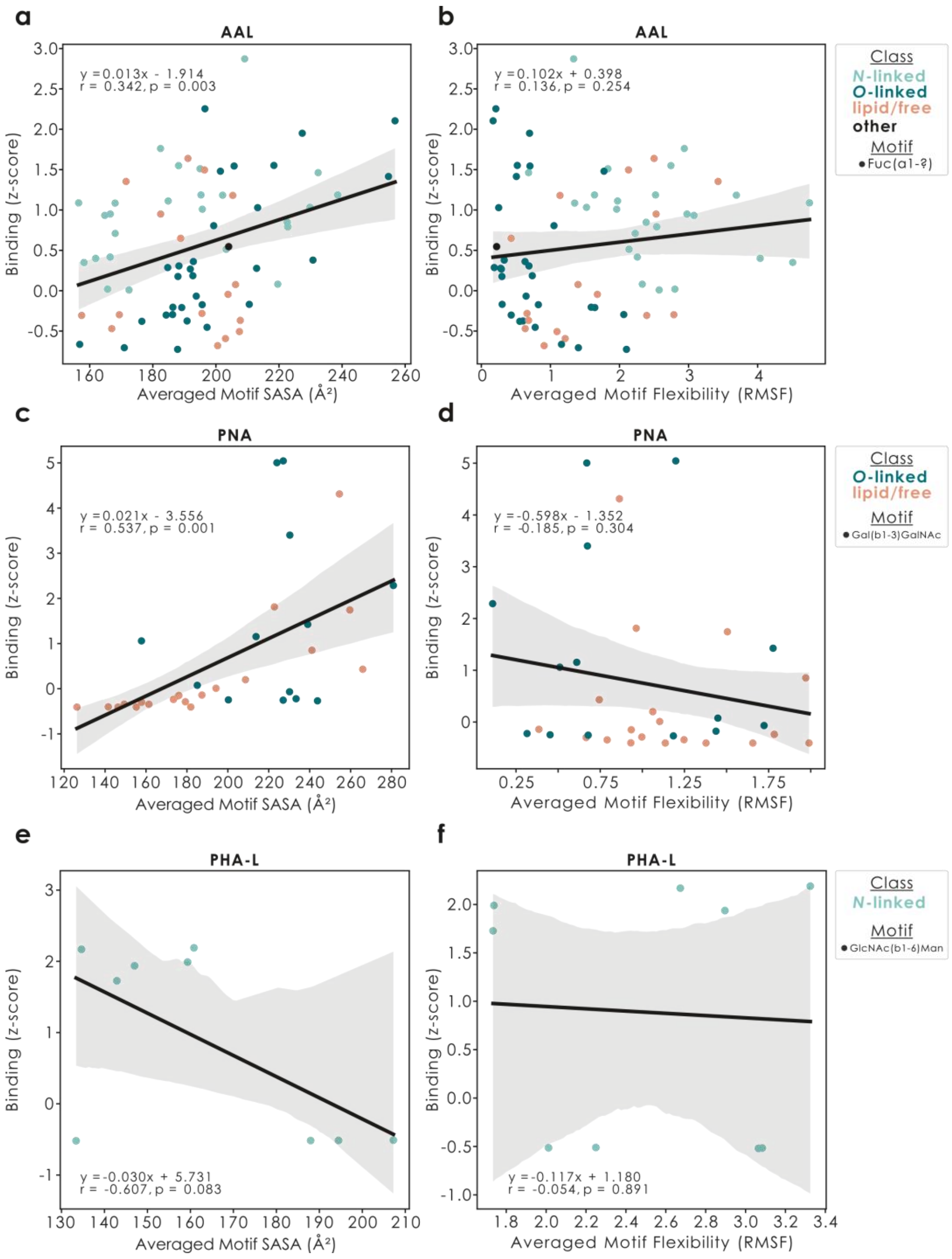


**Supplementary Figure 8. Structural motif properties affect lectin binding. a-f)** For the lectins AAL (a-b), PNA (c-d), and PHA-L (e-f), we used the z-score transformed binding data from glycowork (v1.6) and correlated it with either the averaged SASA (a, c, e) or flexibility (b, d, f) of the literature-known binding motif in each glycan that (i) carried the binding motif, (ii) had binding data, and (iii) was deposited on GlycoShape. On top of the data points as a scatter plot (colored by glycan class), we then drew a linear least-squares regression line (including 95% confidence band) as well as the regression equation, the Pearson's correlation coefficient r, and the p-value of a two-sided t-test of the regression coefficient against zero.

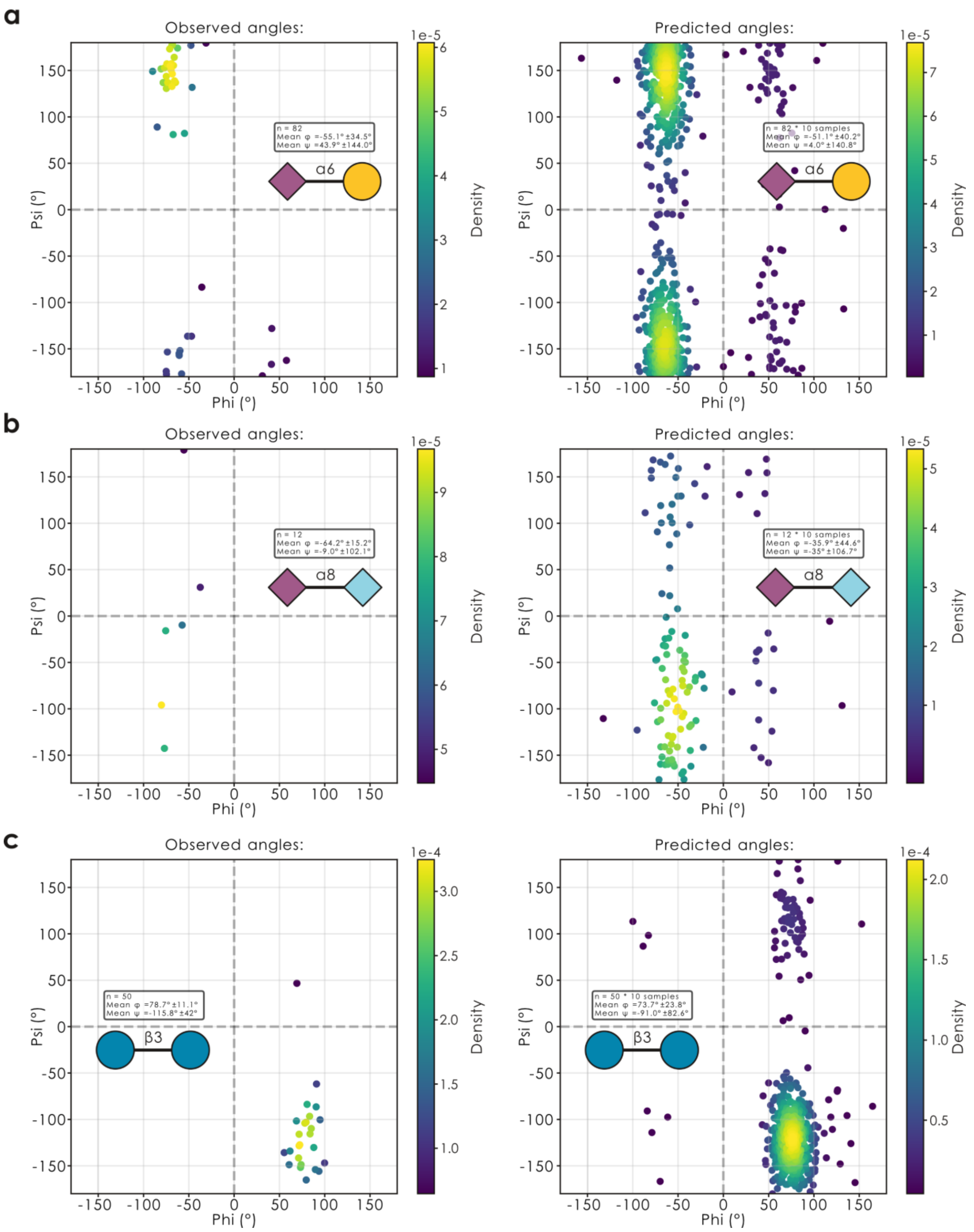


**Supplementary Figure 9. Modeling torsion angles as multimodal von Mises distributions predicts seen and unseen torsion angles accurately. a-c)** For the torsion angles of the disaccharides Neu5Acα2-6Gal (a; seen in training dataset), Neu5Acα2-8Neu5Gc (b; not seen in training dataset), and Glcβ1-3Glc (c; not seen in training dataset), we contrasted observed and predicted torsion angles with the *glycontact.visualize.ramachandran_plot* function. For predicted Ramachandran plots, we used a trained SweetNet-style model by sampling 10 points for each disaccharide sequence occurrence from the multimodal von Mises distribution created from predictions.

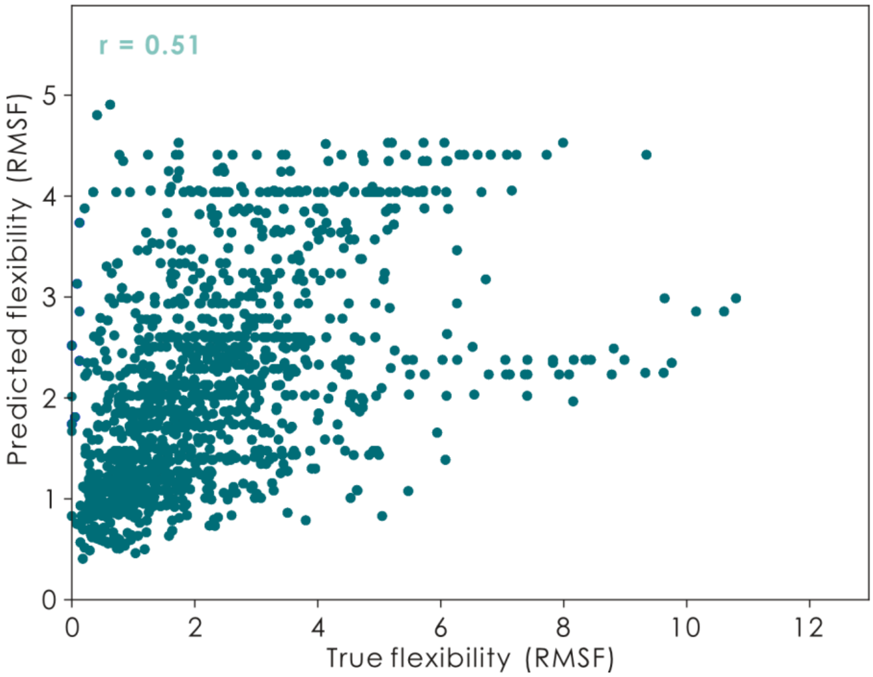


**Supplementary Figure 10. Glycan flexibility predictions correlate with observed flexibilities.** For all glycans in our test set, we extracted their monosaccharide-level flexibility as well as their predicted flexibility using the von Mises-SweetNet model. Shown is a scatter plot of predictions vs ground truths (in root mean square fluctuation or RMSF) as well as their correlation coefficient as Pearson's r.
